# *ViralQuest*: A user-friendly interactive pipeline for viral-sequences analysis and curation

**DOI:** 10.1101/2025.08.10.669577

**Authors:** Gabriel Victor Pina Rodrigues, Lucas Yago Melo Ferreira, Eric Roberto Guimarães Rocha Aguiar

## Abstract

**Background:** High-throughput sequencing (HTS) has become an essential, unbiased tool in virology for identifying known and novel viruses. However, analyzing the large and complex datasets generated by HTS presents significant bioinformatics challenges. The process of accurately identifying and characterizing viral sequences from assembled contigs remains a bottleneck, often requiring specialized expertise and involving non-standardized parameters. There is a pressing need for robust, user-friendly, and reproducible pipelines to streamline this post-assembly analysis.

**Results:** To address these challenges, we developed *ViralQuest*, a bioinformatics tool that automates the in-depth characterization of viral sequences from pre-assembled contigs. The pipeline integrates multiple lines of evidence for robust identification, using Diamond BLASTx against the Viral RefSeq database and pyHMMER searches against the RVDB, Vfam, and eggNOG profile HMM databases. For detailed characterization, *ViralQuest* performs taxonomic classification based on the ICTV nomenclature and functional annotation via Pfam domain analysis.

Novel features of *ViralQuest* include an AI-powered summarization module that uses a Large Language Model (LLM) to generate contextual narratives for key viral findings and a comprehensive confidence score to rank putative viral contigs. All results are consolidated into a single, interactive HTML report that includes dynamic visualizations of contigs, ORFs, and protein domains, alongside detailed data tables that are exportable in TSV and SVG formats.

**Conclusion:** *ViralQuest* provides an accessible and comprehensive solution for the post-assembly analysis of viral metagenomic data. By combining rigorous bioinformatics methods with novel AI-driven features and an intuitive reporting interface, it streamlines the complex process of viral identification and characterization. The tool enhances the interpretability and reliability of results, making in-depth virome analysis more accessible to the broader research community. *ViralQuest* is available on GitHub at https://github.com/gabrielvpina/viralquest/.

## BACKGROUND

The robust detection of viral agents is crucial across diverse scientific domains, from safeguarding public health and ensuring the integrity of biological products to mitigating substantial economic losses in agriculture. High-throughput sequencing (HTS), also referred to as next-generation sequencing (NGS), has emerged as a pivotal technology, fundamentally reshaping the landscape of virology [1]. Its power lies in an unbiased approach, capable of identifying both known and entirely novel viruses without prior sequence knowledge, a clear contrast to traditional targeted methods like PCR that require a priori information about the pathogen. This broad-spectrum capability, not reliant on previous assumptions, is critical for addressing emerging threats and characterizing unknown etiological agents, as evidenced by instances where NGS successfully identified contaminants missed by conventional assays [2].

Despite the transformative capabilities of HTS, the subsequent bioinformatic analysis of the generated voluminous and complex datasets presents considerable challenges that can hinder efficient and accurate virus identification and characterization. While upstream processes generate vast quantities of sequence reads, often dominated by host or non-target microbial nucleic acids, the crucial downstream steps of analyzing assembled contigs for viral signatures, performing reliable taxonomic assignment, and detailed functional annotation remain intricate [3]. The great diversity of viruses, including highly divergent or low-abundance pathogens, demands sophisticated strategies to distinguish true viral sequences and characterize them comprehensively. Furthermore, the current landscape features many analytical tools, often requiring extensive bioinformatics expertise or utilizing subjectively chosen parameters, highlighting an ongoing need for more standardized, robust, reproducible, and user-friendly approaches for in-depth viral characterization from assembled sequence data.

To address these analytical complexities in the post-assembly phase of viral metagenomic studies, we introduce *ViralQuest*, a bioinformatics tool designed to streamline the identification and in-depth characterization of viral sequences from preassembled contigs. *ViralQuest* initiates its workflow by taking FASTA-formatted contigs, employing Open Reading Frame prediction alongside rigorous sequence similarity searches against the Viral RefSeq database and searches against specialized viral profile HMM databases (RVDB, VFam, and eggNOG) to comprehensively identify potential viral conserved regions. This multi-faceted approach enhances the probability of detecting a broad range of viral agents.

Following identification, *ViralQuest* performs extensive characterization of the putative viral contigs. This includes taxonomic classification based on alignment with nt/nr databases and cross-referencing with the ICTV viral taxonomy database, providing assignments from Kingdom to Family. Furthermore, a conserved domain analysis is conducted against the Pfam database to elucidate potential functional elements within the viral sequences. A particularly novel feature of *ViralQuest* is the AI-powered summarization module, which leverages a large language model (local LLM via Ollama or API-based) to interrogate the ICTV Taxonomy Report Database and over 200 viral families’ metadata (including hosts, distribution, and genome characteristics) to generate contextual summaries. The entire analysis culminates in a user-friendly HTML report, integrating all findings—from initial numeric and visual reports regarding sequence identification, structural annotation and taxonomic data to AI-generated summaries— thereby offering researchers a comprehensive and interpretable overview of the viral content within their assembled sequence datasets. Finally, we propose an easy-to-interpret score based on different viral characteristics and sequence annotations to help users to identify high-confident sequences of viral origin.

## METHODS

*ViralQuest* can be installed from GitHub (github.com/gabrielvpina/viralquest/), all instructions and required packages are available in the documentation. The whole pipeline is built in Python and dependencies rely on PyPI packages [4] and Anaconda (Bioconda) packages [5], with one-command installation (Figure 1).

**Figure 1.**
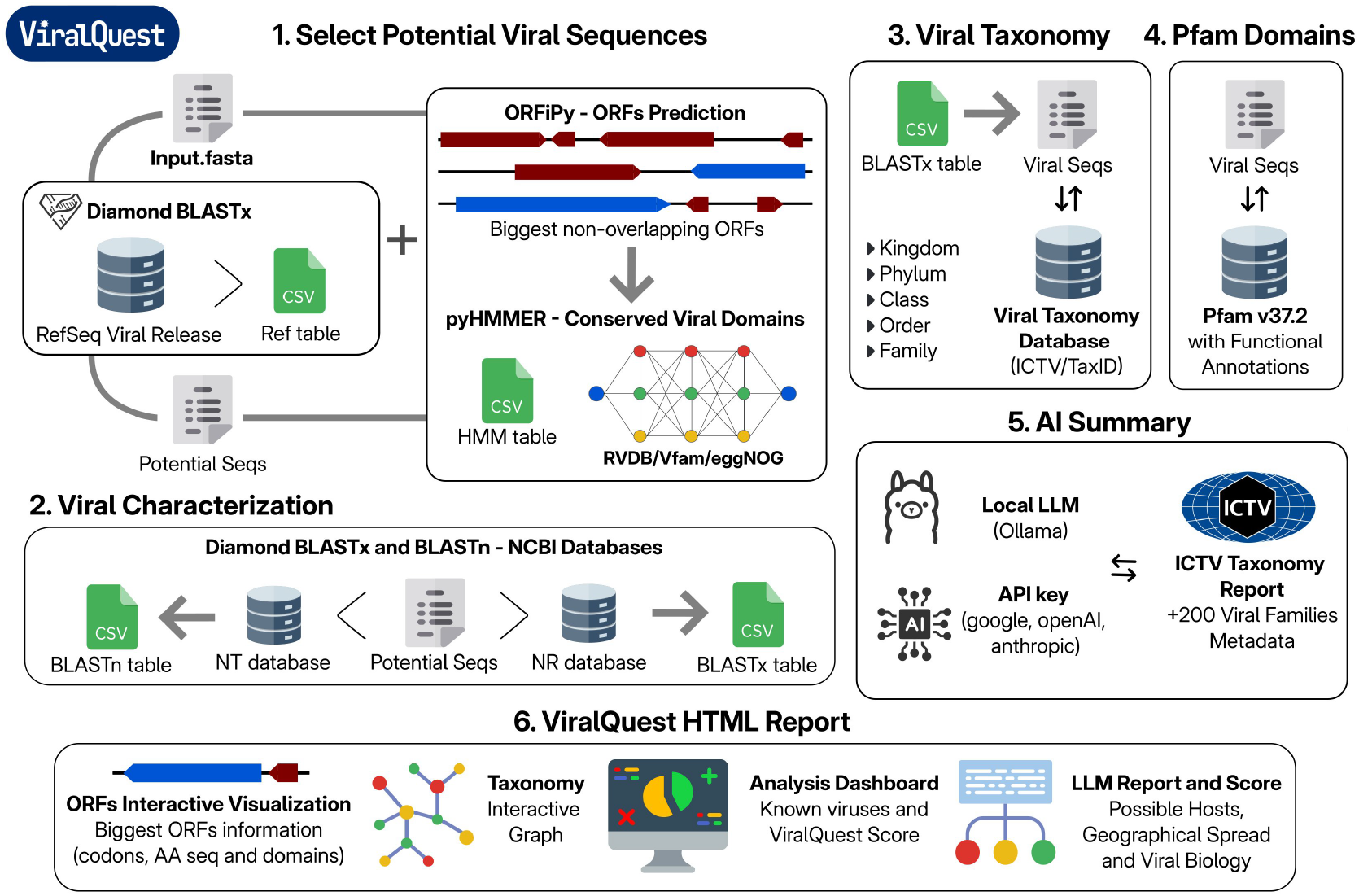
Overview of *ViralQuest* workflow. The pipeline processes input FASTA files through six main stages: (1) Putative viral sequences are identified by alignment against the RefSeq Viral database and by predicting open reading frames (ORFs) containing conserved viral domains from HMM models. (2) These sequences undergo viral characterization via global alignment with the NCBI nucleotide (nt) and non-redundant protein (nr) databases. (3) The taxonomic classification of BLASTx-aligned viral sequences is determined using the ICTV Taxonomy Database. (4) Conserved protein domains are characterized using the Pfam HMM model library. (5) An AI-powered summary integrates the BLAST alignment results, HMM domain information, and ICTV Family Report data. (6) A final HTML report is generated, providing a concise summary of the entire analysis and characterization process.

The *ViralQuest* pipeline is designed for the comprehensive identification and characterization of viral sequences from pre-assembled contigs. The workflow initiates with user-provided sequences in FASTA format and proceeds through several analytical modules to generate a detailed report (**Figure 1**).

### Input Processing and Initial Viral Sequence Identification

Input sequences, assumed to be assembled contigs, are first processed to identify potential protein-coding regions. Open Reading Frames (ORFs) are predicted using ORFiPy [6] and processed to select only the biggest non-overlap ORFs in each contig. Subsequently, to identify sequences of putative viral origin, two parallel homology search strategies are employed. Firstly, the predicted ORFs and the original contig sequences are searched against the NCBI Viral RefSeq protein database using Diamond BLASTx (v2.1.9 or later) [7] with an E-value cutoff of 1×10−5. Secondly, profile Hidden Markov Model (HMM) searches are conducted using pyHMMER (v3.3.2 or later) [8] against several curated viral profile HMM databases: RVDB-prot (v29.8) [9], Vfam (release 228) [10], and eggNOG (v4.5, targeting viral NOGs) [11]. Sequences yielding significant hits through either Diamond BLASTx or pyHMMER searches are selected as candidate viral sequences for downstream characterization.

### Taxonomic and Functional Characterization

Candidate viral sequences undergo detailed taxonomic classification and functional annotation. For taxonomic assignment, sequences are subjected to similarity searches using Diamond BLASTx against the NCBI non-redundant protein (nr) database and BLASTn [12] against the NCBI non-redundant nucleotide (nt) database (including the core_nt subset if specified by the user), employing global alignment strategies. The taxonomic lineage (Kingdom, Phylum, Class, Order, Family) for each viral contig is then determined by cross-referencing the best hits with the NCBI taxonomy database and the International Committee on Taxonomy of Viruses (ICTV) master species list [13].

Functional characterization involves the identification of conserved protein domains. The translated ORFs from candidate viral sequences are scanned against the Pfam-A database (v37.2) [14] using pyHMMER. Significant domain hits provide insights into the potential functions of the encoded viral proteins.

### AI-Powered Analysis Summarization

To provide a contextual overview of the identified viruses, *ViralQuest* incorporates an AI Analysis Summarization module. This module utilizes a Large Language Model (LLM), which can be configured to run locally via Ollama engine [15] or accessed through an API (requiring a user-provided API key). The LLM generates a summary for significant viral findings by integrating information from the ICTV Taxonomy Report Database, which includes detailed descriptions of viral taxa, and a curated knowledge base containing metadata (e.g., host range, geographic distribution, genome characteristics) for over 200 recognized viral families. This aims to provide a concise, informative narrative accompanying the quantitative results.

### Confidence Score

To prioritize and rank the identified putative viral contigs, we developed a comprehensive scoring system (detailed in the **Supplementary file 1**). This system evaluates contigs based on a combination of homology and protein domain evidence. We implemented two parallel scoring approaches: a flexible, criteria-based method using a Large Language Model (LLM), and a deterministic method using a custom Python script, in case of an absent LLM analysis.

Our primary scoring method employed a LLM to assess and assign a score to each contig based on a set of weighted criteria. This approach allowed for a nuanced evaluation of the evidence supporting the viral origin of a sequence. The final score is a composite of three key components, totaling 100 points:

- BLASTn Analysis (30%): This component assesses the nucleotide similarity of the contig to known sequences. The score is determined by both the percentage identity and coverage of the alignment. For instance, a high-confidence hit with over 90% identity and 70% coverage would receive between 25 and 30 points. Lower-quality hits or alignments to non-viral sequences receive progressively fewer points.
- BLASTx Analysis (30%): This evaluates the similarity of the translated nucleotide sequence to known protein sequences. Similar to the BLASTn score, this is weighted by both identity and coverage. A contig with a translated query that aligns to a viral protein with greater than 90% identity and 70% coverage would be awarded between 25 and 30 points.
- HMM Domain Detection (40%): The presence of conserved viral protein domains, as identified by pyHMMER, is a strong indicator of a true viral sequence. This component carries the highest weight in our scoring system. Contigs containing highly significant viral domains (pyHMMER score > 100) are awarded 30 to 40 points, while those with moderately significant or no detectable viral domains receive a lower score.

If any LLM wasn’t provided to execute the pipeline, we provide a stringent rule-based evaluation to implement a secondary scoring system using custom parameters. The script calculates a score based on a series of conditional statements that evaluate the results from the BLASTn and BLASTx analyses. The Python-based score is calculated as follows:

- BLASTx Score: The script first evaluates the BLASTx identity and coverage, assigning up to 25 points for each. The number of points awarded decreases in a stepwise fashion as the identity and coverage values decrease. For example, an identity or coverage greater than 80% contributes 25 points, while a value between 40% and 50% contributes only 5 points.
- BLASTn Score: The evaluation of the BLASTn results is contingent on the presence of a hit. If no BLASTn hit is found, no points are awarded for this category. If a hit is present, 10 points are immediately added. An additional 10 points are awarded if the subject title of the hit contains viral-associated keywords such as “virus” or “phage.” Subsequently, the BLASTn identity and coverage are scored in a manner like the BLASTx analysis, with each contributing a maximum of 15 points.

### Reporting

To facilitate the review and interpretation of bioinformatics results, we developed a custom interactive HTML report using the D3.js JavaScript library [16]. The report dynamically generates a series of visualizations for each putative viral contig. A central feature is a scaled, linear representation of the contig, where predicted ORFs are rendered as arrow-shaped glyphs indicating their position, length, and strand. The fill color of each ORF signifies its status (complete, 5’-partial, or 3’-partial) for rapid assessment. Functionally important regions were highlighted by mapping identified Pfam domains as colored, rounded rectangles beneath their corresponding ORFs, with nucleotide positions calculated based on the parent ORF’s frame and strand. The entire visualization is interactive; hovering over elements reveals tooltips with detailed metadata, including ORF coordinates, sequence data, and functional annotations from BLAST, HMMer, and Pfam searches. Accompanying each contig map are summary panels for BLASTx/n results, taxonomic lineage, and a qualitative AI-generated summary. The report’s dashboard provides a high-level overview, including a dashboard of sequence classification counts and a network graph that visualizes the taxonomic hierarchy (phylum, family, and virus) of the identified sequences.

## FEATURES

### HTML *ViralQuest* Report

The *ViralQuest* pipeline generates a single, interactive HTML file as its final output, providing a comprehensive report for straightforward data exploration. The report is structured into two primary components: an Analysis Summary and In-depth Contig Characterization.

The Analysis Summary offers a high-level overview of the results, featuring:

- A detailed list of contigs identified as viral (**Figure 2B**).
- A taxonomic breakdown of the identified viruses by Order and Family, determined by the best BLAST hit at the amino acid level (**Figure 2C**).
- A visual classification of all processed contigs **(Figure 2A**).

**Figure 2.**
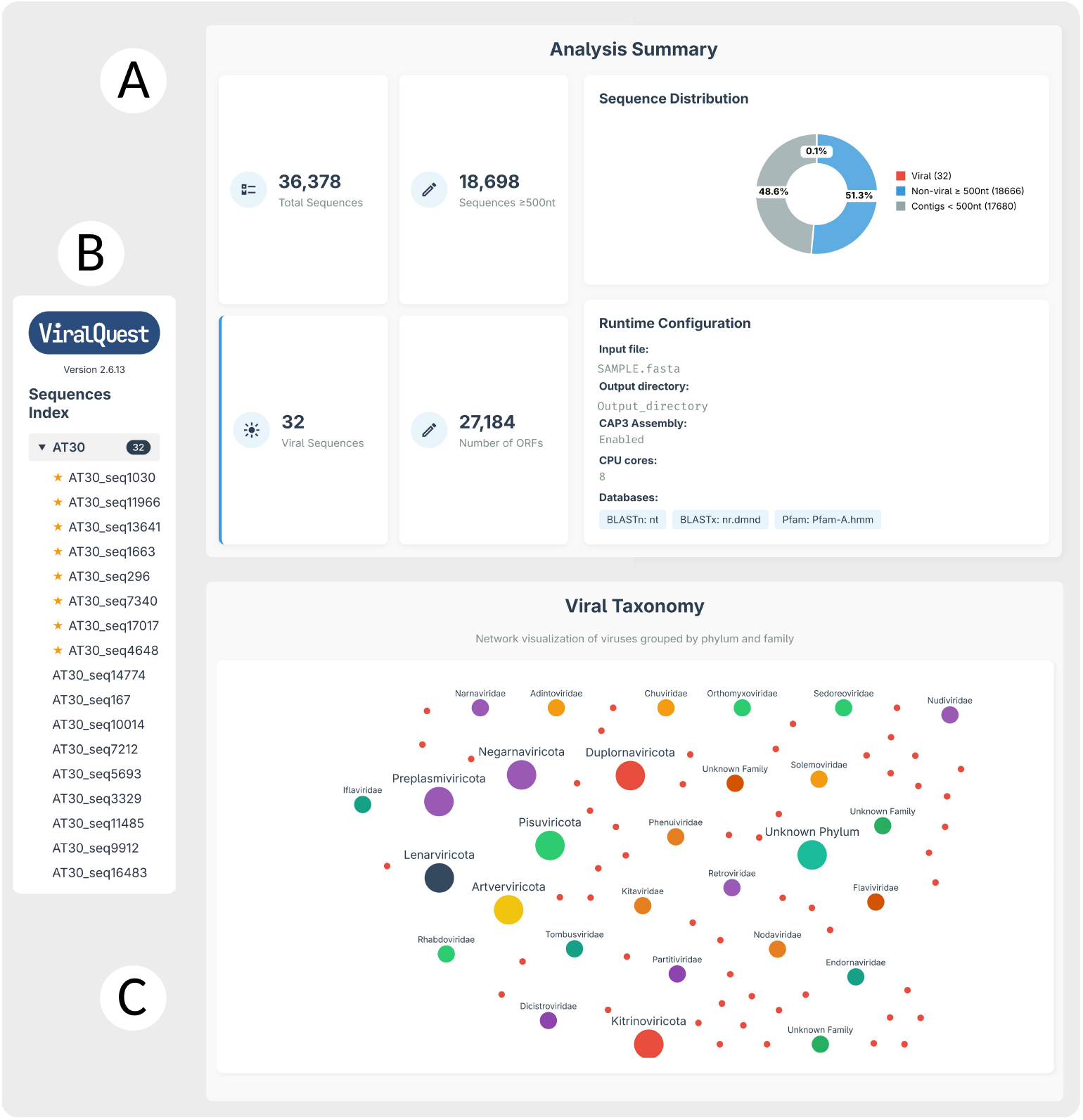
Elements of analysis summary panel in *ViralQuest* HTML report. (A) Dashboard containing general information about sequences in the FASTA input file and an overview of viral sequences distribution. (B) Lateral index of all viral sequences found in the fasta file, named with the prefix of the original FASTA file and the contig number. (C) Taxonomy interactive graph overview with nodes representing Order, Family and Viral species of all found viral sequences. Star in the index panel indicates contig sequences Confidence Score above 80.

The In-depth Contig Characterization section delivers detailed information for each putative viral contig, which includes:

- A header displaying the contig identifier, its length, the top hit from protein similarity searches, and a calculated confidence score (**Figure 3 - Panel 2A**).
- Structural annotation indicating the presence and genomic location of Open Reading Frames (ORFs). Interactive tooltips provide detailed results from pyH- MMER searches against the RVDB, Vfam, eggNOG, and Pfam databases, displaying the accession codes and descriptions of any identified conserved domains (**Figure 3 - Panel 2B**).
- Results from sequence similarity searches, which present detailed metrics from Diamond BLASTx and BLASTn, including the top hit, E-values, percent identity, and alignment coverage (**Figure 3 - Panel 2C and D**).
- Hierarchical taxonomic classification, wherein each viral sequence is classified according to the ICTV nomenclature based on the best hit from the amino acid similarity search (**Figure 3 - Panel 2E**).
- A summary of identified conserved domains based on annotations from the Pfam database (**Figure 3 - Panel 2F**).
- An AI-powered summary of key viral findings, which facilitates the straightforward interpretation of complex viral characterization data (**Figure 3 - Panel 2G**).

**Figure 3.**
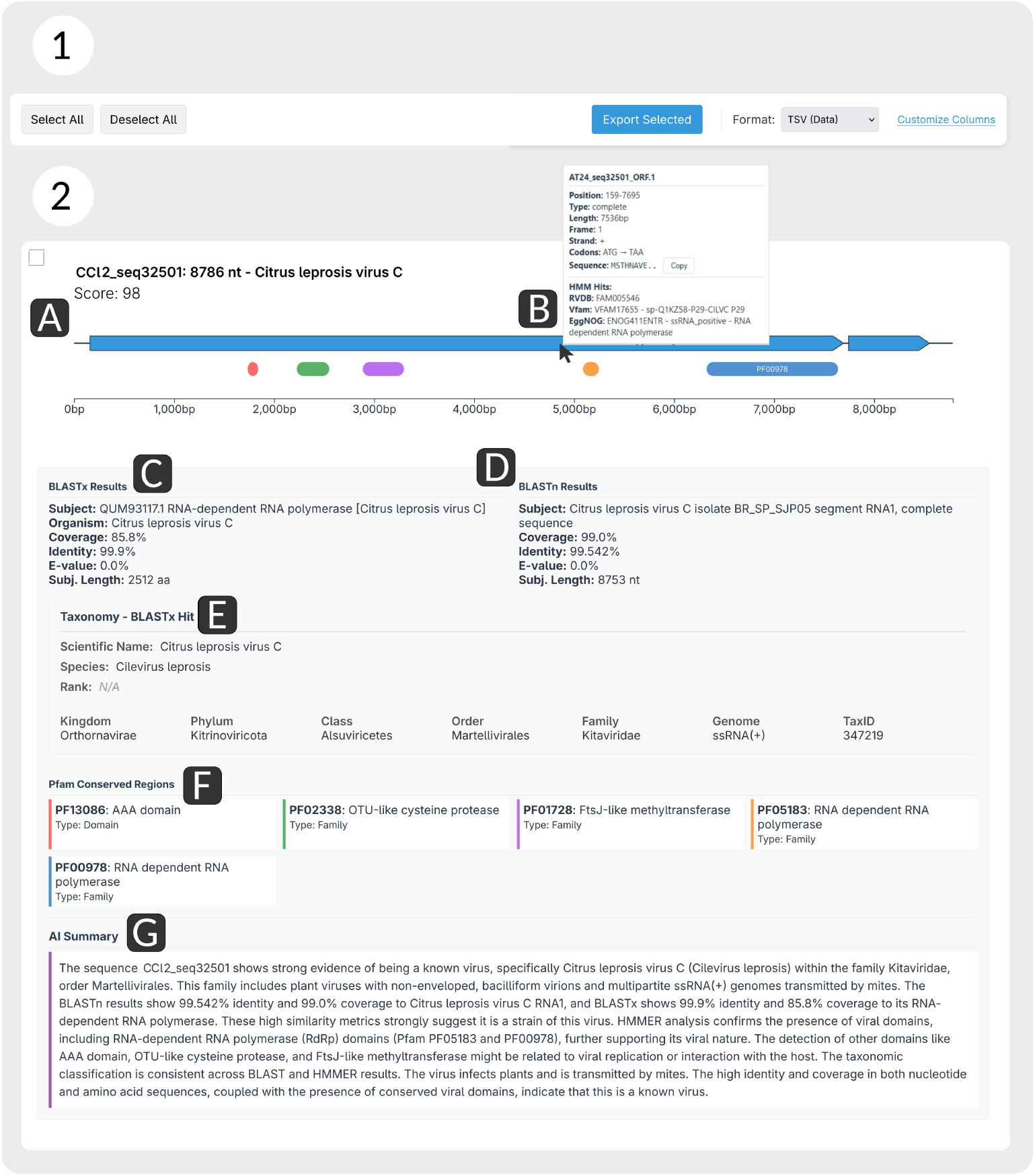
Panel representing the In-depth Contig Characterization section of the *ViralQuest* HTML report. **Panel 1**: Export menu section to create a tabular separated values (TSV) table of selected sequences or export the SVG images from structural annotation plot (ORFs and domains). **Panel 2**: Image of a viral container section with the results of all analysis of the pipeline, (A) Structural Annotation of ORFs and Domains, (B) Mouse Hover Tooltip with ORFs description, (C) BLASTx results, (D) BLASTn results,(E)Taxonomy section of BLASTx Organism hit, (F) Pfam Conserved regions panel and (G) LLM Summary with all information descriptions and viral family metadata from ICTV.

For subsequent computational analyses, the report incorporates functionalities to export data into tab-separated values (TSV) tables and to save ORF diagrams in SVG format. In addition, the tool was implemented to allow the exportation of the whole HTML page as a PDF file through the built-in browser print functionality.

## DECLARATIONS

### Ethics approval and consent to participate

Not applicable

### Consent for publication

Not applicable

### Availability of data and materials

All code and documentation are available on GitHub (https://github.com/gabrielvpina/viralquest)

### Authors’ contributions

Conceptualization, E.R.G.R.A.; methodology, G.V.P.R; formal analysis, G.V.P.R.; resources, E.R.G.R.A.; data curation, G.V.P.R.; writing—original draft preparation, L.Y.M.F.; G.V.P.R. and E.R.G.R.A.; writing—review and editing, G.V.P.R., L.Y.M.F.; and E.R.G.R.A.; visualization, G.V.P.R.; supervision, E.R.G.R.A.; project administration, E.R.G.R.A.; funding acquisition, E.R.G.R.A. All authors have read and agreed to the published version of the manuscript.

## Acknowledgements

We thank all the members of the Virus Bioinformatics Laboratory—UESC for the fruitful discussions and the Laboratory of Molecular Biodiversity and Omics—LabiÔmicas for the local infrastructure. The authors would like to acknowledge the assistance of ChatGPT, a language model developed by OpenAI, in the text editing and grammar reviewing of this manuscript.

## REFERENCES

1. Pérez-Losada M, Arenas M, Galán JC, Bracho MA, Hillung J, García-González N, et al. High-throughput sequencing (HTS) for the analysis of viral populations. Infect Genet Evol. 2020;80:104208.

2. Thorburn F, Bennett S, Modha S, Murdoch D, Gunson R, Murcia PR. The use of next generation sequencing in the diagnosis and typing of respiratory infections. J Clin Virol. 2015;69:96–100.

3. Fitzpatrick AH, Rupnik A, O’Shea H, Crispie F, Keaveney S, Cotter P. High Throughput Sequencing for the detection and characterization of RNA viruses. Front Microbiol. 2021;12:621719.

4. PyPI · the Python Package Index. PyPI. https://pypi.org/. Accessed 6 Jun 2025.

5. Grüning B, Dale R, Sjödin A, Chapman BA, Rowe J, Tomkins-Tinch CH, et al. Bioconda: sustainable and comprehensive software distribution for the life sciences. Nat Methods. 2018;15:475–6.

6. Singh U, Wurtele ES. orfipy: a fast and flexible tool for extracting ORFs. Bioinformatics. 2021;37:3019–20.

7. Buchfink B, Reuter K, Drost H-G. Sensitive protein alignments at tree-of-life scale using DIAMOND. Nat Methods. 2021;18:366–8.

8. Larralde M, Zeller G. PyHMMER: a Python library binding to HMMER for efficient sequence analysis. Bioinformatics. 2023;39.

9. Bigot T, Temmam S, Pérot P, Eloit M. RVDB-prot, a reference viral protein database and its HMM profiles. F1000Res. 2019;8:530.

10. Skewes-Cox P, Sharpton TJ, Pollard KS, DeRisi JL. Profile hidden Markov models for the detection of viruses within metagenomic sequence data. PLoS One. 2014;9:e105067.

11. Huerta-Cepas J, Szklarczyk D, Heller D, Hernández-Plaza A, Forslund SK, Cook H, et al. eggNOG 5.0: a hierarchical, functionally and phylogenetically annotated orthology resource based on 5090 organisms and 2502 viruses. Nucleic Acids Res. 2019;47:D309–14.

12. Camacho C, Coulouris G, Avagyan V, Ma N, Papadopoulos J, Bealer K, et al. BLAST+: architecture and applications. BMC Bioinformatics. 2009;10:421.

13. Master species lists (MSL). Ictv.global. https://ictv.global/msl. Accessed 6 Jun 2025.

14. Sonnhammer EL, Eddy SR, Birney E, Bateman A, Durbin R. Pfam: multiple sequence alignments and HMM-profiles of protein domains. Nucleic Acids Res. 1998;26:320–2.

15. Ollama. Ollama.com. https://ollama.com/. Accessed 6 Jun 2025.

16. Bostock M, Ogievetsky V, Heer J. D3 Data-Driven Documents. IEEE Trans Vis Comput Graph. 2011;17:2301–9.

