## Supplementary File 1 for "*ViralQuest*: A user-friendly interactive pipeline for viral-sequences analysis and curation"

### Overview

This document provides a detailed breakdown of the parameters used to calculate the ViralQuest score (VQ Score) for each contig within our bioinformatics pipeline. We employed two distinct methodologies for scoring: a qualitative assessment guided by a Large Language Model (LLM) and a quantitative, deterministic score calculated via a custom Python script. The parameters for both methods are detailed below.

#### 1 LLM-Based Scoring Parameters

The LLM-based score provides a holistic assessment of viral potential based on weighted criteria totaling 100 points. The model was instructed to assign scores based on the guidelines presented in Table 1.

Table 1: Scoring Criteria for the LLM-Based Methodology.

| Component (Weight) | Criteria Definition | Points Allotted |
|---|---|---|
| **BLASTn Identity & Coverage (30%)** | High Confidence: > 90% identity AND > 70% coverage | 25 – 30 |
| | Medium Confidence: 70-90% identity AND > 50% coverage | 15 – 25 |
| | Low Confidence: < 70% identity OR < 50% coverage | 8 – 15 |
| | No significant data or non-viral hit | 0 – 8 |
| **BLASTx Identity & Coverage (30%)** | High Confidence: > 90% identity AND > 70% coverage | 25 – 30 |
| | Medium Confidence: 70-90% identity AND > 50% coverage | 15 – 25 |
| | Low Confidence: < 70% identity OR < 50% coverage | 0 – 15 |
| **HMM Domain Detection (40%)** | Strong Viral Domains (e.g., HMM score ¿100) | 30 – 40 |
| | Moderate Viral Domains | 20 – 30 |
| | Weak or No Detectable Viral Domains | 0 – 20 |

#### 2 Python-Based Scoring Parameters

A deterministic score was calculated using a custom Python script. This score is a sum of points awarded based on fixed thresholds for BLASTx and BLASTn results. The specific point allocations for each parameter are detailed in Tables 2 and 3.

Table 2: Scoring Rules for BLASTx Results in the Python Script.

| Parameter | Value Range | Points Awarded |
|---|---|---|
| **BLASTx Identity (%)** | $> 80$ | 25 |
| | $70 < \text{value} \leq 80$ | 20 |
| | $60 < \text{value} \leq 70$ | 15 |
| | $50 < \text{value} \leq 60$ | 10 |
| | $40 < \text{value} \leq 50$ | 5 |
| | $\leq 40$ | 1 |
| **BLASTx Coverage (%)** | $> 80$ | 25 |
| | $70 < \text{value} \leq 80$ | 20 |
| | $60 < \text{value} \leq 70$ | 15 |
| | $50 < \text{value} \leq 60$ | 10 |
| | $40 < \text{value} \leq 50$ | 5 |
| | $\leq 40$ | 1 |

Table 3: Scoring Rules for BLASTn Results in the Python Script.

| Parameter | Condition | Points Awarded |
| --- | --- | --- |
| **Base Score** | If a BLASTn hit exists | 10 |
| | If subject title contains 'virus' or 'phage' | +10 (cumulative) |
| **BLASTn Identity (%)** | $> 80$ | 15 |
| | $70 < \text{value} \leq 80$ | 12 |
| | $60 < \text{value} \leq 70$ | 10 |
| | $50 < \text{value} \leq 60$ | 8 |
| | $40 < \text{value} \leq 50$ | 5 |
| | $\leq 40$ | 1 |
| **BLASTn Coverage (%)** | $> 80$ | 15 |
| | $70 < \text{value} \leq 80$ | 12 |
| | $60 < \text{value} \leq 70$ | 10 |
| | $50 < \text{value} \leq 60$ | 8 |
| | $40 < \text{value} \leq 50$ | 5 |
| | $\leq 40$ | 1 |
